# Iron Metabolism and Antioxidant Defense: The Role of Low Intensity-rTMS in Cerebral Ischemia-Reperfusion

**DOI:** 10.1101/2025.09.16.676684

**Authors:** Shumei Fang, Chun Huang, Huimin Wu, Yingmei Xu, Yutong Yao, Chang Yang

## Abstract

**BACKGROUND:** Treating cerebral ischemia/reperfusion injury (CIRI) is challenging, with iron metabolism imbalances and oxidative stress worsening the condition. While low-intensity repetitive transcranial magnetic stimulation (LI-rTMS) shows potential in brain protection, its effects on iron processing and antioxidant use are unclear. This study explored LI-rTMS’s role in regulating iron homeostasis and antioxidant defense for potential therapeutic use in CIRI.

**METHODS:** A rat model of CIRI was established using a filament insertion technique, followed by neurological and infarct volume assessments. In PC12 cells, oxidative damage was induced with H_2_O_2_ or ferric ammonium citrate (FAC), then treated with LI-rTMS or deferoxamine (DFP). Parameters measured included oxidative stress markers, iron metabolism markers, and mitochondrial integrity. Gene knockdown and overexpression experiments explored mechanisms involving LI-rTMS and ACSL4.

**RESULTS:** LI-rTMS improved neurological deficits by boosting GPX4 activity and inhibiting the ACSL4-LPCAT3-LOX axis, reducing lipid peroxidation. It restored iron balance by lowering TFR and DMT1, decreasing ferritin and hepcidin, and increasing FPN for better iron efflux. LI-rTMS also prevented the Fenton reaction, reduced ROS production, and maintained mitochondrial structure and transmembrane potential.

**CONCLUSIONS:** LI-rTMS offers neuroprotection by regulating iron homeostasis through dual pathways: boosting export (via FPN) and reducing uptake/storage (via TFR/DMT1/FE). It also provides ACSL4-dependent antioxidant benefits, suggesting its potential to prevent ferroptosis in CIRI.

## Introduction

Repetitive transcranial magnetic stimulation (rTMS) is a non-invasive, painless therapy that activates cortical neurons using precisely timed, time-varying magnetic fields.^1–4^ The rTMS coil also delivers weaker and diffuse subthreshold stimulation (i.e. low intensity - rTMS) to surrounding brain regions (perifocal stimulation).^5^ Low-intensity repetitive transcranial magnetic stimulation (LI-rTMS), a specific modality of TMS, features the application of relatively weak magnetic fields. While it does not directly induce neuronal action potentials^11^, it indirectly modulates brain function by regulating neuronal excitability, neuroplasticity, and neuronal viability.^6,7^ LI-rTMS demonstrates significantly enhanced safety compared to other TMS forms. As it does not generate high-intensity magnetic pulses, it prevents adverse events such as tinnitus commonly associated with high-intensity TMS.^8^ These properties confer considerable clinical potential in domains such as neurological rehabilitation and psychiatric treatment. Within neurological rehabilitation, LI-rTMS demonstrates efficacy in improving neural deficits and promoting functional recovery for conditions including motor dysfunction, dysphagia, and aphasia—particularly following stroke, spinal cord injury, or peripheral nerve injury. ^5,7,9,10^

Cerebral ischemia/reperfusion injury (CIRI) refers to secondary damage resulting from restored blood flow after arterial occlusion in ischemic stroke. The pathogenesis involves activation of inflammatory responses, excitotoxicity of amino acids, intracellular calcium overload, and induction of oxidative stress.^11,12^ Disruption of cerebral iron homeostasis critically influences secondary damage following ischemic stroke. Recent studies demonstrate that ischemic stroke induces dysregulated iron metabolism in the brain, precipitating pathological iron accumulation in affected regions^13^. The possible cause is an imbalance between the absorption and release of iron,^14^ which is regulated by proteins involved in iron metabolism.^15–17^ Some researches confirm that elevated ferrous ions from intracellular iron overload trigger pathological accumulation of reactive oxygen species (ROS) via Fenton reactions, subsequently driving lipid peroxide generation and ferroptosis.^18,19^ Interestingly, our investigation demonstrated that LI-TMS confers beneficial antioxidant effects in treating CIRI in rats.^20^ Thus, this study plans to further illustrate the mechanism underlying its amelioration of brain ischemia-reperfusion injury. We believe that LI-rTMS may delay intracellular iron accumulation by modulating iron-regulatory proteins. This intervention could consequently reduce ROS levels and enhance neurological recovery following cerebral ischemia-reperfusion injury.

To validate the above hypothesis, rat CIRI models were generated via the intraluminal filament method. Neuroprotective effects of LI-rTMS were evaluated using macro-scale indices including neurological deficit scores, cerebral infarction volume, and oxidative stress biomarkers. For mechanistic investigation, *in vitro* PC12 cell models simulating oxidative stress and iron overload conditions were established. Ferric ammonium citrate (FAC) induced iron overload, hydrogen peroxide (H₂O₂) triggered oxidative stress states, and deferiprone (DFP) served as an iron-chelating control to identify key targets of LI-rTMS. Integrated analysis of iron metabolism and oxidative stress indicators elucidated underlying molecular pathways. This study not only confirms LI-rTMS efficacy at the whole-animal level but provides mechanistic evidence for its regulation of iron-redox homeostasis.

## METHODS

Data supporting the present study are available from the corresponding author upon reasonable request. A detailed description of the methods is found in the Supplemental Material.

### Animals

Male Sprague-Dawley rats (SPF-grade, 9–12 weeks old, 260 ± 20 g) were procured from Changsha Tianqing Biotechnology Co., Ltd. (Animal License No. SCXK (Xiang) 2019-0014, China). All experimental protocols received approval from the Animal Ethics Committee of Guizhou Medical University (Approval No. 2304418), complying with China’s Laboratory Animal Management Regulations (Ministry of Health Document No. 55, 2001). Before experimentation, animals underwent a 7-day acclimatization period in a controlled environment (23 ± 2°C, 55 ± 5% relative humidity). No clinical abnormalities were observed during acclimatization.

### Establishment of Rat Ischemia/Reperfusion (I/R) Model

Rats were anesthetized, and the left common carotid artery (CCA) and external carotid artery (ECA) were exposed. A middle cerebral artery occlusion (MCAO) suture (Model 2838-A4; Beijing Xinotian Biotechnology Co., Ltd., China) was advanced 20 mm intracranially through the internal carotid artery (ICA) to occlude the middle cerebral artery (MCA). After 1 h of ischemia, the suture was withdrawn to initiate reperfusion, and the CCA was ligated prior to wound closure. Sham-operated animals underwent identical surgical procedures excluding arterial occlusion.

### Cell Culture Maintenance and Experimental Treatment

PC12 cells (rat pheochromocytoma cell line) were obtained from the Cell Cryopreservation Center of Pharmaceutical Sciences Research Laboratory, Guizhou Medical University. Cells were maintained in DMEM (Gibco, USA) supplemented with 10% fetal bovine serum (FBS, OriCell, China) and 1% penicillin/streptomycin (Solarbio, China) in humidified culture flasks at 37℃ with 5% CO_2_. ACSL4 knockdown and overexpression RNA (Genecefe Biotechnology Corporation, Wuxi, China) were used for transfection. Transduced cells underwent selection with 2μg/mL pyrimethamine (Beyotime Biotechnology, China) to establish stable knockdown (KD) and over-expression (OE) cell lines. PC12 cells were exposed to 1200 μM hydrogen peroxide (H₂O₂; Tianjin Damao Chemical Co., Ltd., China) for 30 min to induce oxidative stress. Cells were treated with 4 mM ferrous amino citrate (FAC; Aladdin, Shanghai, China) for 24 h to prepare an iron overload model, and 50 mM deferoxamine (DFP; Ferriprox®, Apotex Inc., Canada) was used for 24 h as a positive control. LI-rTMS was applied using a portable transcranial magnetic stimulator (Shijiazhuang Dukang Medical Devices, China) with parameters: 50 Hz frequency, intensity level 2 (6-9 mT), and 30 min duration per session.

### Data Analysis

All statistical analyses were performed using SPSS 26.0. For comparisons among multiple groups, a one-way analysis of variance (ANOVA) was conducted, followed by post hoc tests (LSD or Dunnett’s T3, as appropriate). Values are expressed as mean ± SD (n ≥ 3). Statistical significance was set at *P* < 0.05.

### Results

#### LI-rTMS Demonstrates Neuroprotective Benefits against CIRI

The research created a MCAO model in adult SD rats via intraluminal filament occlusion, followed by interventions LI-rTMS, DFP, and FAC (Figures 1A). Compared to sham-operated controls, both the MCAO model and FAC groups exhibited significantly elevated mNSS scores (Figure 1B), confirming successful model establishment. Concurrently, postoperative body weight monitoring revealed significant weight loss in the MCAO and FAC groups (Figure 1C), indicative of metabolic disturbances. Notably, LI-rTMS and DFP interventions significantly reduced neurological deficit scores and attenuated weight loss (Figure 1B and 1C), demonstrating their neuroprotective effects. Tissue viability was further assessed using TTC staining. As shown in Figures 1D and 1E, MCAO and FAC groups displayed markedly increased infarct volumes compared with sham controls. Both LI-rTMS and DFP treatments substantially reduced infarct areas, with LI-rTMS exhibiting superior therapeutic efficacy. In summary, LI-rTMS treatment significantly promoted the recovery of neurological function, reduced pathological weight loss, and effectively reduced the area of cerebral infarction.

**Figure 1.**
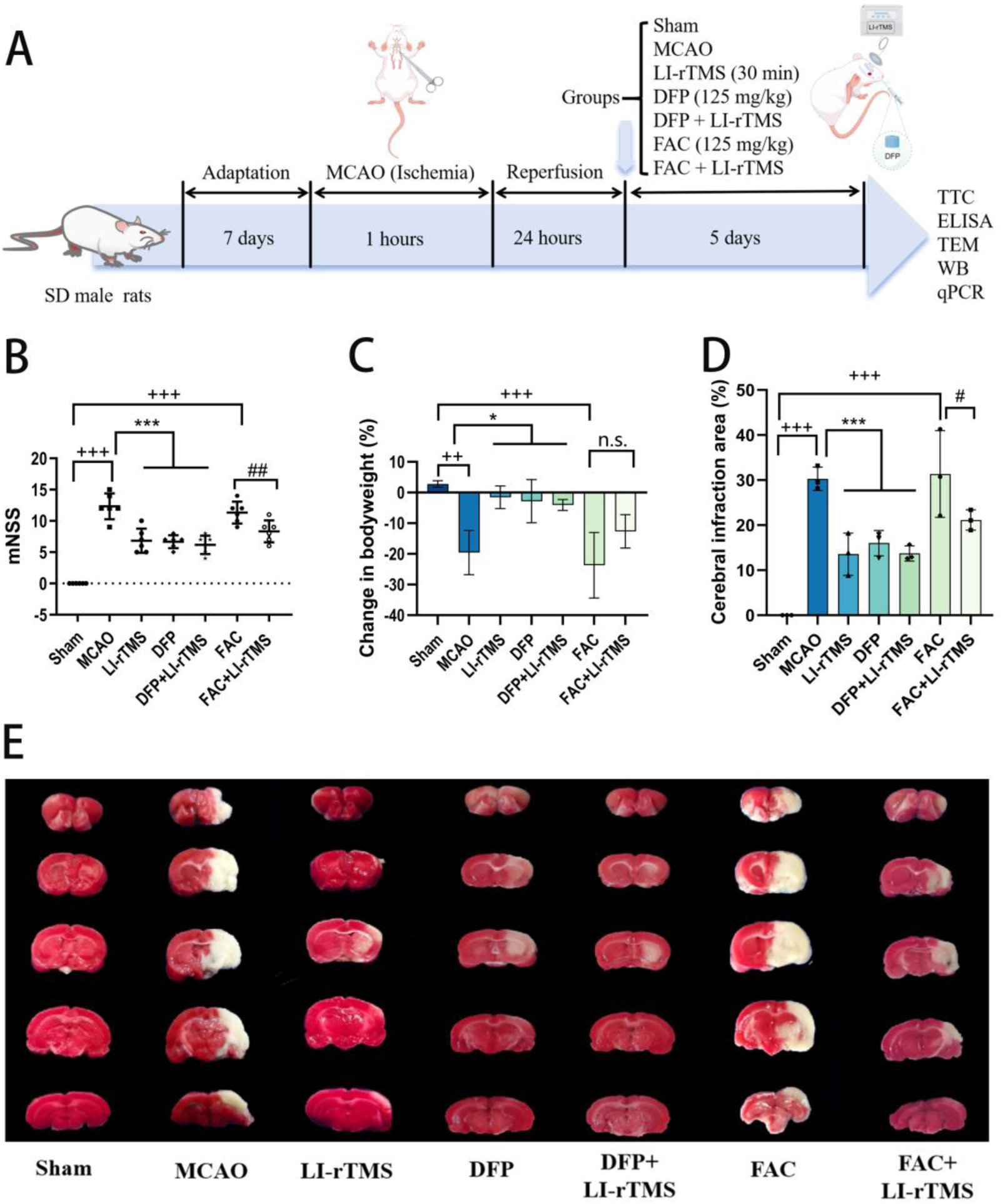
Neuroprotective effects of LI-rTMS in a rat MCAO model. **A**, Schematic of experimental design demonstrating stroke model induction and therapeutic interventions. **B**, Neurological score (mNSS) 24 hours after surgery. n=6. **C**, Percent body weight change in rats after 5 days of treatment. n=6. **D**, Quantitative infarct volume analysis. **E**, Representative TTC-stained brain sections. Scale bar = 5 mm. Results expressed as mean ± SD; significance by ANOVA: ^++^*P* < 0.01, ^+++^*P* < 0.001 vs. sham group; \*\*\**P* < 0.05, \**P* < 0.001 vs. I/R (MCAO) group; ^#^*P* < 0.05, ^##^*P* < 0.01 vs. FAC group; n.s., not significant.

### LI-rTMS Attenuates Oxidative Stress Induced by CIRI

Previous studies have demonstrated that LI-rTMS exhibits antioxidative properties, evidenced by significantly diminished lipid peroxidation (malondialdehyde, MDA) and elevated superoxide dismutase (SOD) activity, protecting rats against CIRI.^20^ To elucidate the mechanisms of LI-rTMS in reducing oxidative stress, an *in vitro* oxidative stress model (Figure 2A) was developed using hydrogen peroxide (H₂O₂; 1200 μM, 30 min), in parallel with an iron overload model induced by ferric ammonium citrate (FAC; 4 mM, 24 h). Optimal treatment parameters—deferiprone (DFP; 50 μM, 24 h) concentration and LI-rTMS duration (30 min)—were subsequently determined (Figure S1). BODIPY581/591 C11 probing revealed significantly increased LPO levels following H₂O₂/FAC exposure (Figure 2B and 2C). Comparable results were observed for ROS via DCFH-DA fluorescent detection (Figure 2D). Both LI-rTMS and DFP ameliorated oxidative stress in H₂O₂ models; however, LI-rTMS failed to reduce FAC-induced LPO (Figure 2B and 2C). Antioxidant marker analysis confirmed suppressed GSH/GSSG ratios, NADPH, and GPX4 in H₂O₂/FAC groups. LI-rTMS and DFP raised these markers in H₂O₂/FAC groups (Figure 2E, 2F and 2G).

**Figure 2.**
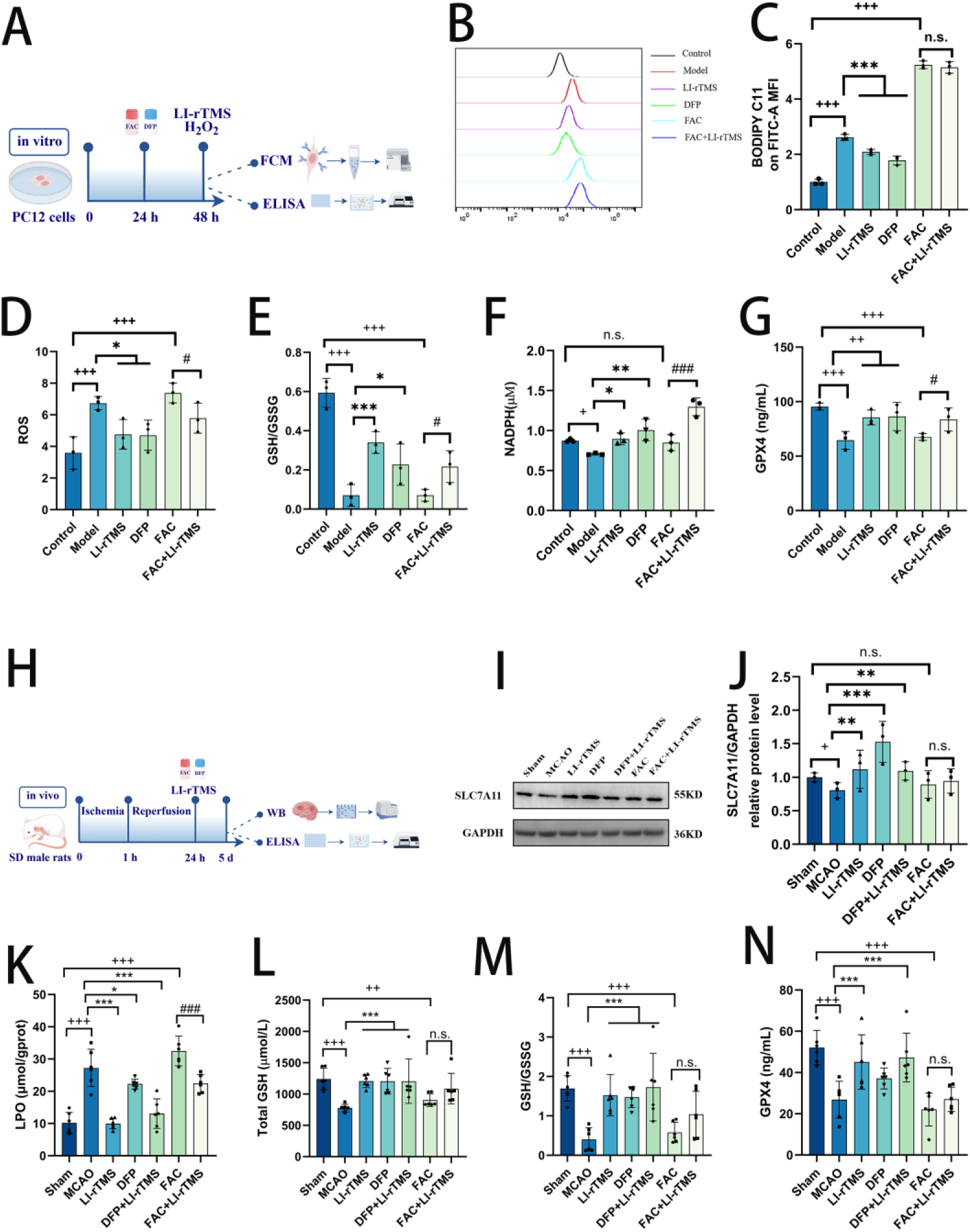
LI-rTMS alleviates CIRI by modulating redox balance. **A-G**, Cellular mechanisms: **A**, Schematic diagram of the *in vitro* model, **B**-**C**, Flow cytometry histograms and mean fluorescence intensity analysis of LPO via BODIPY 581/591 C11. n=3, **D**, ROS fluorescence intensity. n=3, **E**, GSH/GSSG ratio. n=3, **F**, NADPH levels. n=3, **G**, GPX4 enzymatic activity. n=3. **H-L**, Oxidative stress parameters in rat brain tissue: **H**, Timeline of *in vivo* experiments, **I-J**, SLC7A11 protein expression detected by western blotting and quantified via ImageJ. n=3, **K**, LPO levels. n=6, **L**, Total glutathione (GSH) content. n=6, **M**, GSH/GSSG ratio. n=6, **N**, GPX4 enzymatic activity. n=6. Results expressed as mean ± SD; significance by ANOVA: ^++^*P*<0.01, ^+++^*P*<0.001 vs sham group; \**P*<0.05, \*\*\**P*<0.001 vs MCAO (I/R model) group; ^#^*P*<0.05, ^##^*P*<0.01 vs FAC group; n.s., not significant.

To systematically investigate LI-rTMS neuroprotective mechanisms, oxidative stress markers and key antioxidant components were evaluated *in vivo* (Figure 2H). Quantitative analyses indicated elevated LPO levels in MCAO and FAC groups compared with sham controls, suggesting aggravated oxidative damage. Notably, LI-rTMS and DFP treatment effectively reduced LPO levels, with LI-rTMS demonstrating superior efficacy (Figure 2K). Western blotting analyses revealed significantly downregulated SLC7A11 expression in MCAO groups, which was restored by LI-rTMS intervention. However, FAC induction did not alter SLC7A11 expression, and LI-rTMS lacked upregulatory effects (Figure 2I and 2J). Total glutathione quantification showed markedly reduced GSH content in MCAO rats versus controls, restored by LI-rTMS (Figure 2L). Measurements of GSH/GSSG ratios confirmed LI-rTMS-mediated restoration of redox equilibrium (Figure 2M). Furthermore, GPX4 expression was substantially downregulated in MCAO groups, while both LI-rTMS and DFP significantly enhanced GPX4 levels (Figure 2N). However, FAC models LI-rTMS showed no efficacy in GSH/GPX4 modulation.

Collectively, LI-rTMS confers neuroprotection via synergistic mechanisms, including enhanced SLC7A11/GSH-mediated antioxidant defenses, upregulated GPX4 expression, and effective ROS scavenging, thereby attenuating ischemia/reperfusion– induced oxidative injury.

### LI-rTMS can Significantly Decrease Cellular Fe^2+^ Levels in CIRI

To elucidate the potential antioxidant mechanism of LI-rTMS and its association with iron metabolism, this study further validated the iron-regulatory effects of LI-rTMS in cellular models through analysis of iron-related parameters and regulatory proteins. Results demonstrated significant pathological Fe²⁺ accumulation in cells under oxidative stress and iron overload conditions, LI-rTMS significantly attenuated this pathological accumulation (Figure 3A). ELISA demonstrated significant upregulation of TFR, DMT1, FE, and hepcidin expression coupled with downregulation of FPN in H₂O₂/FAC groups. Crucially, LI-rTMS treatment reversed pathological iron accumulation by suppressing TFR/DMT1/FE/hepcidin expression while enhancing FPN expression (Figure 3B, 3C, 3D, 3E and 3F).

**Figure 3.**
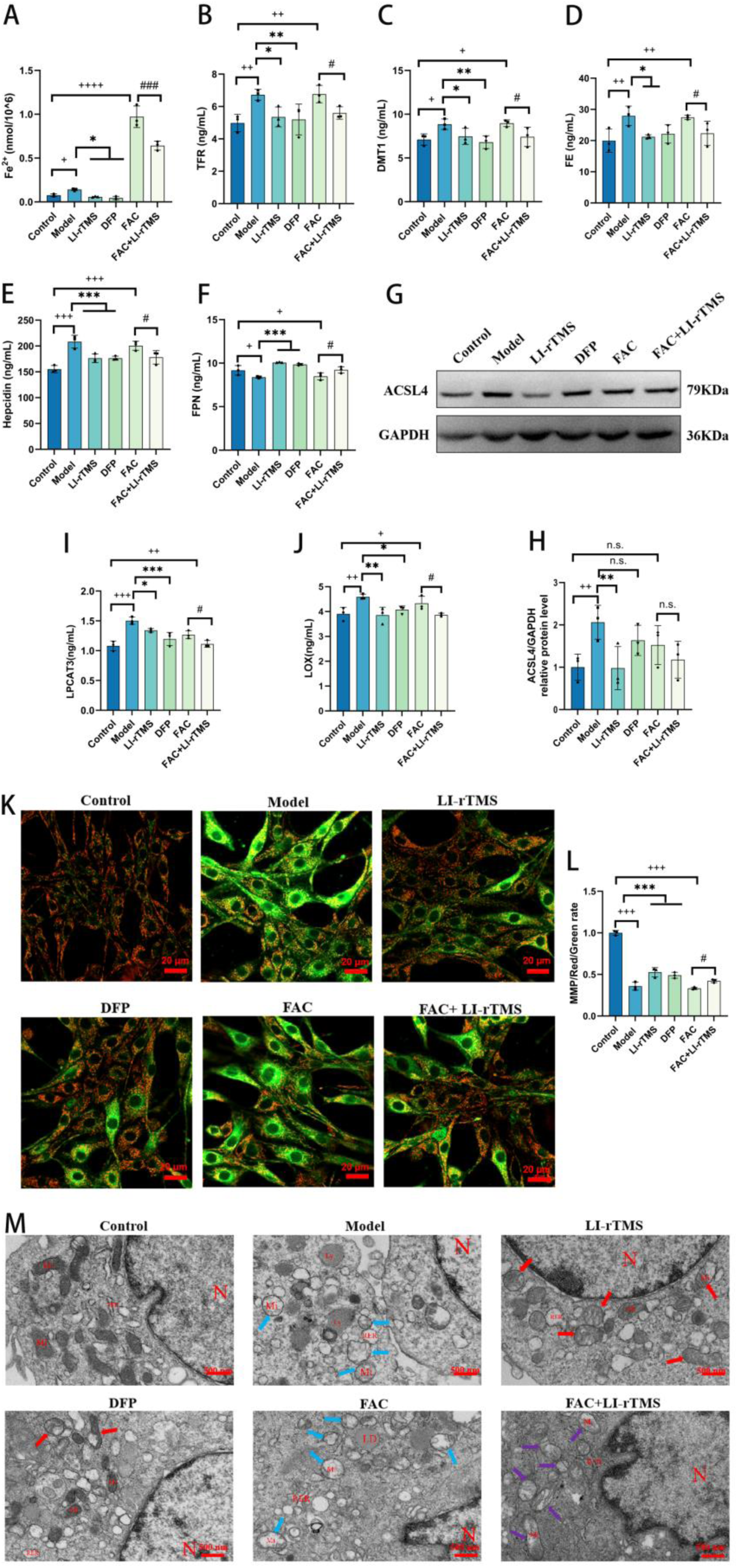
Regulatory effects of LI-rTMS on iron homeostasis and mitochondrial function in an oxidative stress and iron overload PC12 cell model. **A-F**, Iron metabolism analysis: **A**, Fe²⁺ levels in PC12 cells. n=3, **B**, TFR expression. n=3, **C**, DMT1 expression. n=3, **D**, FE expression. n=3, **E**, hepcidin expression. n=3, **F**, FPN expression. n=3. **G-J**, Lipid metabolism protein analysis: **G**, ACSL4 expression, **H**, Quantified ACSL4 protein expression. n=3, **I**, LPCAT3 expression. n=3, **J**, LOX expression levels. n=3. **K-M**, Mitochondrial protection: **K**, Mitochondrial membrane potential (ΔΨm) detected by JC-1 reagent. n=3. **L**, quantitative analysis of ΔΨm, **M**, TEM revealed disrupted mitochondrial cristae (blue arrows) in I/R-injured neurons, with structural improvement following LI-rTMS treatment (scale bar: 500 nm). Data are presented as mean ± SD (n=3).vs. control group: ^+++^*P* < 0.001, ^++^*P* < 0.01, ^+^*P* < 0.05;vs. model group: \*\*\**P* < 0.001, \*\**P* < 0.01, \**P* < 0.05;vs. FAC group: ^#^*P* < 0.05.n.s., not statistically significant.

Western blotting analysis indicated significantly elevated ACSL4 protein levels in model groups versus sham controls. LI-rTMS intervention normalized ACSL4 expression, whereas other treatment groups showed no significant alterations (Figure 3G and 3H). Moreover, LI-rTMS attenuated oxidative stress and iron overload-induced increases in LPCAT3 and LOX expression (Figure 3I and 3J), suggesting potential lipid metabolism regulation through these proteins.

TEM showed significant mitochondrial damage in model and FAC groups, marked by ruptured membranes and disrupted cristae. Both LI-rTMS and DFP treatment significantly restored mitochondrial architecture, with improved membrane integrity and cristae morphology (Figure 3K and 3L). Mitochondrial membrane potential assays demonstrated significant depolarization in H₂O₂/FAC groups, which was substantially reversed by LI-rTMS and DFP administration (Figure 3M), indicating mitochondrial functional preservation.

Furthermore, iron metabolism analysis *in vivo* revealed pathological iron dysregulation patterns consistent with *in vitro* findings: both MCAO and FAC-induced iron overload models exhibited significant iron homeostasis perturbations compared to sham-operated controls. Specifically, cerebral Fe²⁺ levels were markedly elevated, whereas serum total iron content and TIBC decreased significantly. Although altered transferrin saturation (TSAT) was observed, it showed no statistically significant difference across treatment groups (Figure 4D), collectively indicating pathological iron accumulation (Figure 4A, 4B, 4C, and 4D). LI-rTMS ameliorated these abnormalities in MCAO groups but failed to regulate Fe²⁺ and TIBC levels in FAC-induced models (Figure 4B and 4C). This differential efficacy suggests FAC-mediated pathological iron overload may disrupt fundamental regulatory pathways (e.g., redox-sensitive iron transporters), limiting LI-rTMS’s neuromodulatory capacity — a finding consistent with its inability to modulate GSH/GPX4 axes previously identified.

**Figure 4.**
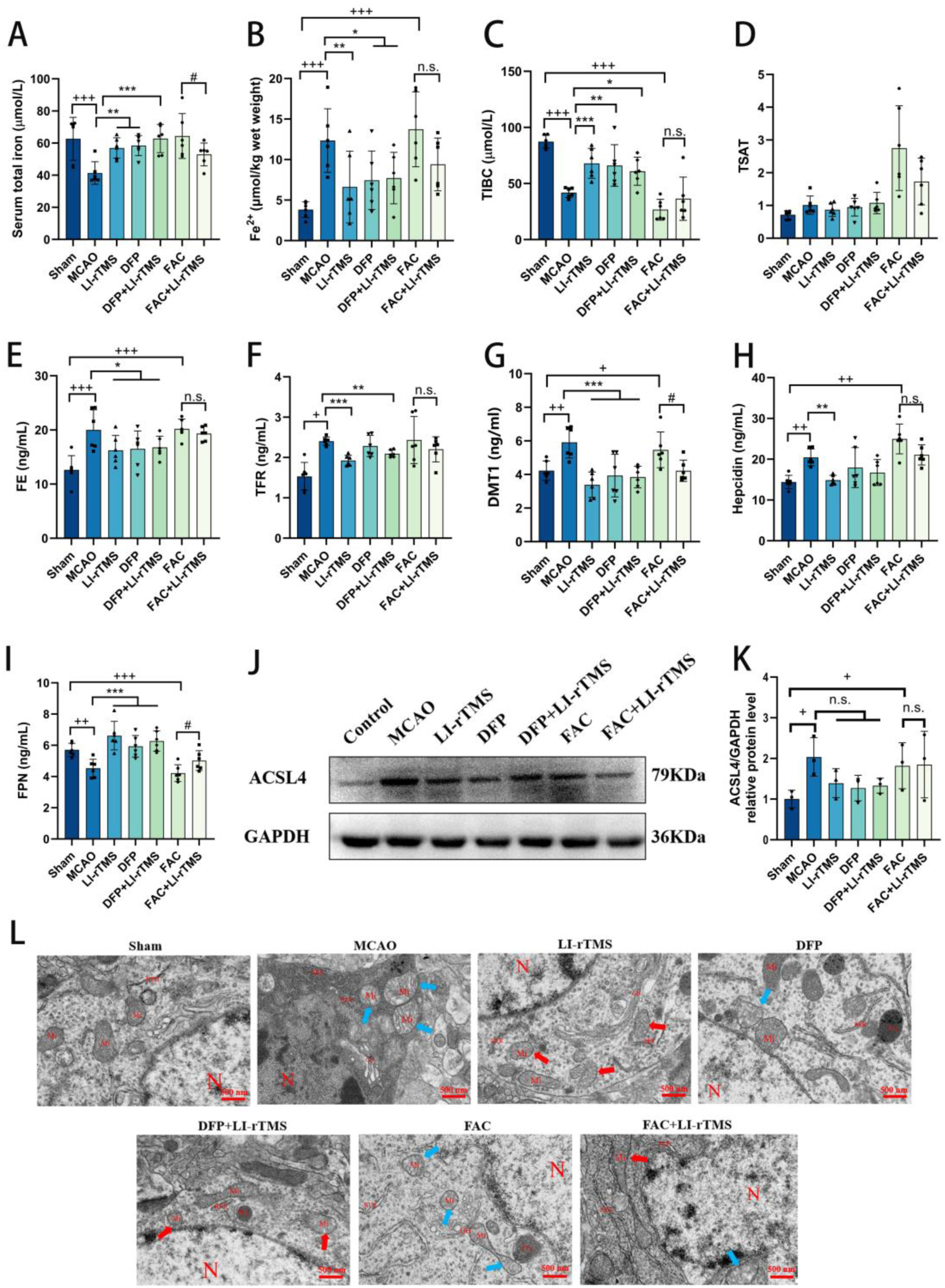
LI-rTMS can significantly decrease cellular Fe^2+^ levels *in vivo*. **A-D**, Iron metabolism analysis. n=6: **A**, Serum total iron levels, **B**, Cerebral Fe²⁺ content, **C**, Total iron-binding capacity (TIBC), **D**, Transferrin saturation (TSAT). **E-I**, Expression of iron metabolism-related proteins in brain tissue (ELISA). n=6: **E**, FE, **F**, TFR, **G**, DMT1, **H**, Hepcidin, **I**, FPN. **J**, Western blotting analysis of ACSL4 protein expression. **K**, Quantification of ACSL4 protein expression. n=3. **L**, TEM revealing mitochondrial cristae disruption in ischemia-reperfusion injured neurons (blue arrowhead) and structural restoration post LI-rTMS treatment (Scale bar: 500 nm). Results expressed as mean ± SD; significance by ANOVA: vs. Sham group: ^++^*P* < 0.01, ^+++^*P* < 0.001;vs. I/R model (MCAO) group: \**P* < 0.05, \*\*\**P* < 0.001;vs. FAC group: ^#^*P* < 0.05, ^##^*P* < 0.01;n.s., not statistically significant.

ELISA revealed concomitant alterations in cerebral iron-regulatory proteins, including upregulated TFR and DMT1 (implicating iron uptake), elevated ferritin and hepcidin (suggesting iron storage and regulation), and downregulated FPN (suppressing iron efflux) (Figure 4E, 4F,4G, 4H, and 4I). LI-rTMS treatment restored these proteins in MCAO groups; however, it failed to significantly downregulate TFR, ferritin and hepcidin in FAC cohorts (Figure 4E,4F and 4H). As anticipated from disrupted iron homeostasis, FAC-induced perturbations were less responsive to intervention. MCAO groups exhibited significantly elevated ACSL4 protein expression, unaltered by LI-rTMS or DFP treatment (Figure 4J and 4K). TEM demonstrated severe mitochondrial damage in MCAO and FAC groups, featuring membrane disruption, cristae disorganization, and matrix dissolution (Figure 4L). TEM demonstrated severe mitochondrial pathological alterations in MCAO and FAC groups, including membrane disruption, cristae disorganization, and matrix dissolution (Figure 4L). LI-rTMS administration substantially restored mitochondrial architecture and improved membrane integrity and cristae morphology.

Collectively, these findings demonstrate that CIRI may induce iron dysregulation and lipid peroxidation. LI-rTMS alleviates pathological iron accumulation by downregulating TFR and DMT1 to reduce serum ferritin and hepcidin concentrations, while upregulating FPN to enhance iron efflux capacity. Moreover, it inhibits the ACSL4-LPCAT3-LOX signaling axis, exhibiting potent antioxidant activity. This neuroprotective mechanism preserves mitochondrial ultrastructural integrity and membrane potential (ΔΨm).

### Modulating ACSL4 in Ferroptosis: Transcriptional-Translational Discrepancy Underlying LI-rTMS Efficacy

Previous studies indicate that LI-rTMS exerts neuroprotection by suppressing iron-dependent lipid peroxidation. To dissect its molecular targets, we investigated ACSL4 – a key enzyme in ferroptosis execution – using ACSL4-overexpressing (OE) and ACSL4-knockdown (KD) PC12 cell models (Figure S6). qPCR confirmed significant downregulation of ACSL4 mRNA by LI-rTMS in all genetic backgrounds, including KD cells (Figure 5A). Notably, while LI-rTMS decreased ACSL4 protein in WT and OE cells, it failed to further suppress ACSL4 protein in KD cells (Figure 5B and 5C). This aligns with evidence that ACSL4 activity is essential for maintaining basal membrane phospholipid remodeling; its expression may be constrained near a physiological threshold to ensure cell survival, thereby limiting further downregulation. This threshold phenomenon resolves the apparent paradox between mRNA suppression and protein stability in KD cells.

**Figure 5.**
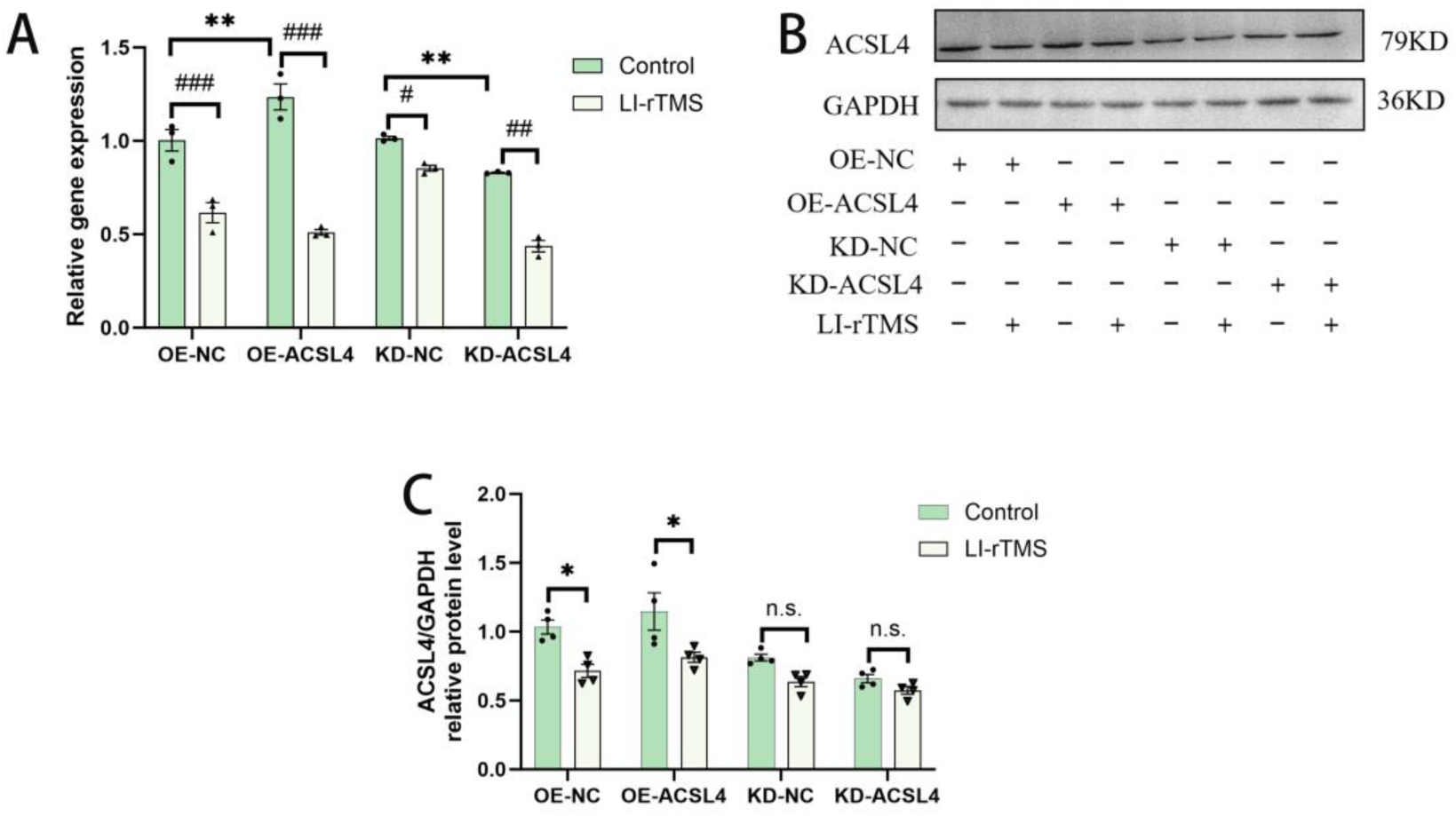
Validation of genetic models and LI-rTMS-mediated ACSL4 modulation in ferroptosis: **A**, ACSL4 transcriptional regulation by LI-rTMS in OE/KD PC12 cells. n=3 (\**P*<0.05, \*\**P*<0.01 vs. OE-NC/KD-NC; ^#^*P*<0.05, ^##^*P*<0.01, ^###^*P*<0.001 vs. Control). **B-C**, LI-rTMS effects on ACSL4 protein expression in OE/KD PC12 models. n=4 (\**P*<0.05 vs. Control).

## Discussion

Globally, stroke remains a significant health issue, responsible for 11.6% of disability-adjusted life years lost to non-communicable diseases. It is the leading cause of early death in those over 60, affecting quality of life and life expectancy.^21^ Ischemic stroke, caused by cerebral artery blockage, makes up 70-80% of cases, with around 69.9 million incidents worldwide in 2021.^22^ Current treatments include tissue plasminogen activator (tPA), effective in 31-50% of cases,^23^ and endovascular thrombectomy (EVT), which has an 88% success rate for large vessel blockages.^24^

However, these treatments are limited by strict time windows (tPA within 4.5 hours, EVT within 6 hours) and risk of reperfusion injury.^25^ Cerebral ischemia/reperfusion injury (CIRI) results from complex mechanisms like oxidative stress, neuroinflammation, and ferroptosis.^26^ While animal studies show that anti-inflammatory agents (such as minocycline, statin therapy, resveratrol) offer neuroprotection, clinical trials reveal limited success and notable side effects, such as hepatotoxicity and gastrointestinal toxicity,^27^ highlighting the urgent need for better treatments.

LI-rTMS has shown promise in preclinical studies for reducing neurological deficits and oxidative stress-related damage in rats with cerebral ischemia.^28–31^ However, the use of low-intensity rTMS (LI-rTMS) is debated due to its weaker magnetic field, which limits motor evoked potentials.^5^ Against this backdrop, the therapeutic potential of LI-rTMS warrants in-depth exploration from a mechanistic perspective.

According to new research, pathological iron buildup occurs in ischemic areas after stroke due to disruption of the blood-brain barrier.^17,32,33^ Our experimental results show significant iron deposition in both oxidative stress-treated PC12 cells (Figure 3A) and the rat CIRI model (Figure 4A and 4B), which is consistent with this mechanism. Subsequent research showed that the expression of proteins linked to iron metabolism, such as FE, DMT1, and TFR, was significantly upregulated under ischemic-hypoxic condition (Figure 3B, 3C, 3D, 3E and 3F).^34,35^ This dysregulation of iron regulatory proteins promotes increased intracellular free iron concentrations, thereby exacerbating cerebral tissue damage. These findings corroborate previous studies indicating that iron metabolism dysfunction represents a critical pathological mechanism in ischemic-hypoxic brain injury. Our findings indicate that LI-rTMS exerts regulatory effects on cerebral iron metabolism during ischemia-reperfusion. Specifically, LI-rTMS treatment demonstrated the capacity to reduce excessive iron accumulation in brain tissue through coordinated modulation of iron transport proteins: (1) upregulation of FPN expression and (2) downregulation of TFR, DMT1, FE, and hepcidin expression (Figure 4E 4F, 4G, 4H and 4I). This multi-target regulation significantly reduces the concentration of free ferrous ions, thereby inhibiting the burst of hydroxyl radicals derived from the Fenton reaction, providing direct experimental evidence for the positive feedback hypothesis linking iron metabolism imbalance and oxidative stress. Under ischemic conditions, cerebral iron accumulation is influenced by systemic iron circulation, wherein circulating Fe³⁺ binds to transferrin (Tf) for subsequent delivery to tissues.^36^ The transferrin saturation (TSAT) index, representing the percentage of iron-bound transferrin in serum, serves as a clinically relevant biomarker of systemic iron status.^37^ While previous studies in stroke models associate elevated TSAT with exacerbated neuronal damage and TSAT reduction with neuroprotection, our results revealed no significant intergroup differences in TSAT levels (Figure 4C and 4D), indicating that the exogenous iron overload model partially blocked the iron-regulatory effects of LI-rTMS, confirming that its mechanism of action lies in the restoration of endogenous iron homeostasis rather than direct iron chelation. Critically, the limited efficacy of LI-rTMS in regulating iron parameters in FAC-treated models (e.g., unchanged Fe²⁺, TIBC, hepcidin; Figures 4B/C/H) reinforces its primary mechanism: targeting redox imbalance rather than direct iron chelation. FAC-induced iron accumulation likely triggers cell death through iron-autonomous pathways (e.g., mitochondrial iron toxicity), which are less dependent on ACSL4-mediated lipid peroxidation –explaining the differential responsiveness to LI-rTMS between ischemia (MCAO) and pure iron-overload (FAC) models.

During cerebral ischemia-reperfusion (I/R) injury in rats, excessive ROS production occurs primarily due to mitochondrial electron transport chain dysfunction or NAD(P)H oxidase overactivation, leading to exacerbated oxidative stress.^18^ Additionally, intracellular Fe²⁺ overload reacts with H₂O₂, generating highly oxidative hydroxyl radicals (·OH) that further induce ferroptosis.^38^ This peroxidative cascade preferentially targets mitochondrial membranes due to their high polyunsaturated fatty acid content, culminating in phospholipid hydroperoxide accumulation that directly compromises membrane integrity. GPX4 is a pivotal enzyme that catalyzes the reduction of lipid hydroperoxides into non-toxic lipid alcohols, thereby mitigating oxidative damage. Its activity is strictly glutathione (GSH)-dependent,^39^ as both GSH and its oxidized form (GSSG) serve as critical antioxidants that protect cells from ROS-mediated injury. Notably, ischemic injury disrupts the equilibrium between system Xc- and glutamate transport. This imbalance subsequently impairs intracellular glutamate uptake and consequently compromises GSH synthesis, exacerbating oxidative stress in the infarcted region.^40^ Damage of this kind weakens the antioxidant defenses of the cell, which raises lipid ROS and causes ferroptosis. The functional subunit of system Xc-, SLC7A11, has a high glutamate and cystine specificity. It facilitates the synthesis of GSH to shield cells from oxidative stress and preserve redox homeostasis by mediating extracellular glutamate release and cystine uptake. This procedure successfully stops lipid peroxidation-induced cell death.^41^ According to our experimental results, rats exposed to CIRI and oxidative stress show markedly elevated ROS levels and lipid peroxidation accumulation, both of which are reduced by LI-rTMS treatment (Figure 2B, 2C, 2D and 2K). Additionally, it has been demonstrated that selenium supplementation increases GPX4 expression, providing stroke-affected rats with neuroprotection against ferroptosis.^42^ Our study demonstrates that LI-rTMS protects against cerebral ischemia-reperfusion injury in rats through three interconnected mechanisms: (1) upregulation of SLC7A11 expression (Figure 2I, and 2J), (2) elevation of the GSH/GSSG ratio (Figure 2M), and (3) enhancement of GPX4 expression with concomitant suppression of lipid peroxidation (Figure 2N). Recent work by Chen et al. demonstrated that ACSL4 suppression attenuates ferroptosis following cerebral ischemia and promotes post-stroke neurological recovery.^43^ Conversely, elevated ACSL4 expression exacerbates ischemic brain injury. Our *in vitro* experiments demonstrated significant upregulation of ACSL4 expression in PC12 cells following oxidative stress. This induction was effectively suppressed by LI-rTMS intervention (Figure 3G and 3H).

In summary, this study elucidate how LI-rTMS antagonizes CIRI through a dual-track mechanism: (1) Iron metabolism level: systemically regulating iron homeostasis (reducing iron input/storage mediated by TFR/DMT1/ferritin and promoting iron output via FPN) significantly lowers free Fe²⁺ concentration to inhibit the Fenton reaction; (2) At the lipid peroxidation level: activation of the SLC7A11-GSH-GPX4 antioxidant axis and inhibition of ACSL4 block the mitochondrial membrane phospholipid peroxidation cascade. The synergistic reduction of ROS-mediated oxidative stress provides an innovative mechanism for the neuroprotective effects of LI-rTMS.

## Conclusion

This study explored the impact of LI-rTMS on iron homeostasis and oxidative stress responses following CIRI. LI-rTMS treatment significantly ameliorated neurological deficits, prevented abnormal body weight loss, and reduced cerebral infarct volume. Mechanistically, LI-rTMS modulated the expression of iron metabolism-related proteins, promoting Fe^2+^ efflux while limiting its cellular uptake. This effect subsequently attenuated Fenton reaction initiation ROS generation. At the molecular level, LI-rTMS restored cerebral iron homeostasis by upregulating GPX4 expression and suppressing the ACSL4-LPCAT3-LOX axis, thereby inhibiting lipid peroxidation. These coordinated mechanisms conferred antioxidant protection, mitigated mitochondrial dysfunction, and ultimately alleviated ischemia/reperfusion-induced brain injury.

## Acknowledgments

Chang Yang designed the experiments and acquired funding. Shumei Fang and Chun Huang performed both cell culture assays and animal experiments. Huimin Wu and Yingmei Xu participated in animal experiments and data collection, while Yutong Yao contributed to data collation and organization. Chang Yang reviewed the article. All authors read and approved the final article.

## Supplemental Material

Supplemental Methods

Tables S1

Figures S1–S7

## References

1. Klomjai W, Katz R, Lackmy-Vallée A. Basic principles of transcranial magnetic stimulation (TMS) and repetitive TMS (rTMS). Ann Phys Rehabil Med. 2015;58:208–213. doi: 10.1016/j.rehab.2015.05.005

2. Lefaucheur JP, Aleman A, Baeken C, Benninger DH, Brunelin J, Di Lazzaro V, Filipović SR, Grefkes C, Hasan A, Hummel FC, et al. Evidence-based guidelines on the therapeutic use of repetitive transcranial magnetic stimulation (rTMS): An update (2014-2018). Clin Neurophysiol. 2020;131:474–528. doi: 10.1016/j.clinph.2019.11.002

3. Du J, Yang F, Hu J, Hu J, Xu Q, Cong N, Zhang Q, Liu L, Mantini D, Zhang Z, et al. Effects of high- and low-frequency repetitive transcranial magnetic stimulation on motor recovery in early stroke patients: Evidence from a randomized controlled trial with clinical, neurophysiological and functional imaging assessments. Neuroimage Clin. 2019;21:101620. doi: 10.1016/j.nicl.2018.101620

4. Xu P, Huang Y, Wang J, An X, Zhang T, Li Y, Zhang J, Wang B. Repetitive transcranial magnetic stimulation as an alternative therapy for stroke with spasticity: a systematic review and meta-analysis. J Neurol. 2021;268:4013–4022. doi: 10.1007/s00415-020-10058-4

5. Moretti J, Rodger J. A little goes a long way: Neurobiological effects of low intensity rTMS and implications for mechanisms of rTMS. Curr Res Neurobiol. 2022;3:100033. doi: 10.1016/j.crneur.2022.100033

6. Makowiecki K, Garrett A, Harvey AR, Rodger J. Low-intensity repetitive transcranial magnetic stimulation requires concurrent visual system activity to modulate visual evoked potentials in adult mice. Sci Rep. 2018;8:5792. doi: 10.1038/s41598-018-23979-y

7. Moretti J, Terstege DJ, Poh EZ, Epp JR, Rodger J. Low intensity repetitive transcranial magnetic stimulation modulates brain-wide functional connectivity to promote anti-correlated c-Fos expression. Sci Rep. 2022;12:20571. doi: 10.1038/s41598-022-24934-8

8. Rossi S, Antal A, Bestmann S, Bikson M, Brewer C, Brockmöller J, Carpenter LL, Cincotta M, Chen R, Daskalakis JD, et al. Safety and recommendations for TMS use in healthy subjects and patient populations, with updates on training, ethical and regulatory issues: Expert Guidelines. Clin Neurophysiol. 2021;132:269–306. doi: 10.1016/j.clinph.2020.10.003

9. Sheng R, Chen C, Chen H, Yu P. Repetitive transcranial magnetic stimulation for stroke rehabilitation: insights into the molecular and cellular mechanisms of neuroinflammation. Front Immunol. 2023;14:1197422. doi: 10.3389/fimmu.2023.1197422

10. Bai G, Jiang L, Ma W, Meng P, Li J, Wang Y, Wang Q. Effect of Low-Frequency rTMS and Intensive Speech Therapy Treatment on Patients With Nonfluent Aphasia After Stroke. Neurologist. 2020;26:6–9. doi: 10.1097/nrl.0000000000000303

11. Jolugbo P, Ariëns RAS. Thrombus Composition and Efficacy of Thrombolysis and Thrombectomy in Acute Ischemic Stroke. Stroke. 2021;52:1131–1142. doi: 10.1161/strokeaha.120.032810

12. Wu MY, Yiang GT, Liao WT, Tsai AP, Cheng YL, Cheng PW, Li CY, Li CJ. Current Mechanistic Concepts in Ischemia and Reperfusion Injury. Cell Physiol Biochem. 2018;46:1650–1667. doi: 10.1159/000489241

13. McCarthy RC, Sosa JC, Gardeck AM, Baez AS, Lee CH, Wessling-Resnick M. Inflammation-induced iron transport and metabolism by brain microglia. J Biol Chem. 2018;293:7853–7863. doi: 10.1074/jbc.RA118.001949

14. Spence H, McNeil CJ, Waiter GD. The impact of brain iron accumulation on cognition: A systematic review. PLoS ONE. 2020;15:e0240697. doi: 10.1371/journal.pone.0240697

15. Rouault TA. Iron metabolism in the CNS: implications for neurodegenerative diseases. Nat Rev Neurosci. 2013;14:551–564. doi: 10.1038/nrn3453

16. Koleini N, Shapiro JS, Geier J, Ardehali H. Ironing out mechanisms of iron homeostasis and disorders of iron deficiency. J Clin Invest. 2021;131. doi: 10.1172/jci148671

17. DeGregorio-Rocasolano N, Martí-Sistac O, Ponce J, Castelló-Ruiz M, Millán M, Guirao V, García-Yébenes I, Salom JB, Ramos-Cabrer P, Alborch E, et al. Iron-loaded transferrin (Tf) is detrimental whereas iron-free Tf confers protection against brain ischemia by modifying blood Tf saturation and subsequent neuronal damage. Redox Biol. 2018;15:143–158. doi: 10.1016/j.redox.2017.11.026

18. Wan J, Ren H, Wang J. Iron toxicity, lipid peroxidation and ferroptosis after intracerebral haemorrhage. Stroke Vasc Neurol. 2019;4:93–95. doi: 10.1136/svn-2018-000205

19. Alim I, Caulfield JT, Chen Y, Swarup V, Geschwind DH, Ivanova E, Seravalli J, Ai Y, Sansing LH, Ste Marie EJ, et al. Selenium Drives a Transcriptional Adaptive Program to Block Ferroptosis and Treat Stroke. Cell. 2019;177:1262–1279.e1225. doi: 10.1016/j.cell.2019.03.032

20. Wang L, Yang S, Li L, Huang Y, Li R, Fang S, Jing J, Yang C. A low-intensity repetitive transcranial magnetic stimulation coupled to magnetic nanoparticles loaded with scutellarin enhances brain protection against cerebral ischemia reperfusion injury. J Drug Delivery Sci Technol. 2022;74:103606. doi: 10.1016/j.jddst.2022.103606

21. Feigin VL, Stark BA, Johnson CO, Roth GA, Bisignano C, Abady GG, Abbasifard M, Abbasi-Kangevari M, Abd-Allah F, Abedi V, et al. Global, regional, and national burden of stroke and its risk factors, 1990–2019: a systematic analysis for the Global Burden of Disease Study 2019. The Lancet Neurology. 2021;20:795–820. doi: 10.1016/S1474-4422(21)00252-0

22. Li XY, Kong XM, Yang CH, Cheng ZF, Lv JJ, Guo H, Liu XH. Global, regional, and national burden of ischemic stroke, 1990-2021: an analysis of data from the global burden of disease study 2021. EClinicalMedicine. 2024;75:102758. doi: 10.1016/j.eclinm.2024.102758

23. Emberson J, Lees KR, Lyden P, Blackwell L, Albers G, Bluhmki E, Brott T, Cohen G, Davis S, Donnan G, et al. Effect of treatment delay, age, and stroke severity on the effects of intravenous thrombolysis with alteplase for acute ischaemic stroke: a meta-analysis of individual patient data from randomised trials. Lancet. 2014;384:1929–1935. doi: 10.1016/s0140-6736(14)60584-5

24. Goyal M, Menon BK, van Zwam WH, Dippel DWJ, Mitchell PJ, Demchuk AM, Dávalos A, Majoie CBLM, van der Lugt A, de Miquel MA, et al. Endovascular thrombectomy after large-vessel ischaemic stroke: a meta-analysis of individual patient data from five randomised trials. The Lancet. 2016;387:1723–1731. doi: 10.1016/S0140-6736(16)00163-X

25. Zhang M, Liu Q, Meng H, Duan H, Liu X, Wu J, Gao F, Wang S, Tan R, Yuan J. Ischemia-reperfusion injury: molecular mechanisms and therapeutic targets. Signal Transduct Target Ther. 2024;9:12. doi: 10.1038/s41392-023-01688-x

26. Zhao K, Wang P, Tang X, Chang N, Shi H, Guo L, Wang B, Yang P, Zhu T, Zhao X. The mechanisms of minocycline in alleviating ischemic stroke damage and cerebral ischemia-reperfusion injury. European Journal of Pharmacology. 2023;955:175903. doi: 10.1016/j.ejphar.2023.175903

27. Mallah K, Couch C, Borucki DM, Toutonji A, Alshareef M, Tomlinson S. Anti-inflammatory and Neuroprotective Agents in Clinical Trials for CNS Disease and Injury: Where Do We Go From Here? Front Immunol. 2020;11:2021. doi: 10.3389/fimmu.2020.02021

28. De Michele M, Piscopo P, Costanzo M, Lorenzano S, Crestini A, Rivabene R, Manzini V, Petraglia L, Iacobucci M, Berto I, et al. Can Repetitive Transcranial Magnetic Stimulation (rTMS) Promote Neurogenesis and Axonogenesis in Subacute Human Ischemic Stroke? Biomedicines. 2024;12. doi: 10.3390/biomedicines12030670

29. Fan S, Yan L, Zhang J, Qian Y, Wang M, Yang L, Yu T. Effects of repetitive transcranial magnetic stimulation on lower extremity motor function and optimal parameters in stroke patients with different stages of stroke: a systematic evaluation and meta-analysis. Front Neurol. 2024;15:1372159. doi: 10.3389/fneur.2024.1372159

30. Wang M, Zhang W, Zang W. Repetitive transcranial magnetic stimulation improves cognition, depression, and walking ability in patients with Parkinson’s disease: a meta-analysis. BMC Neurol. 2024;24:490. doi: 10.1186/s12883-024-03990-9

31. Lu QQ, Zhu PA, Li ZL, Holmes C, Zhong Y, Liu H, Bao X, Xie JY. Efficacy of Repetitive Transcranial Magnetic Stimulation Over the Supplementary Motor Area on Motor Function in Parkinson’s Disease: A Meta-analysis. Am J Phys Med Rehabil. 2025;104:318–324. doi: 10.1097/phm.0000000000002593

32. Tian X, Li X, Pan M, Yang LZ, Li Y, Fang W. Progress of Ferroptosis in Ischemic Stroke and Therapeutic Targets. Cell Mol Neurobiol. 2024;44:25. doi: 10.1007/s10571-024-01457-6

33. Liu X, Chen C, Han D, Zhou W, Cui Y, Tang X, Xiao C, Wang Y, Gao Y. SLC7A11/GPX4 Inactivation-Mediated Ferroptosis Contributes to the Pathogenesis of Triptolide-Induced Cardiotoxicity. Oxid Med Cell Longev. 2022;2022:3192607. doi: 10.1155/2022/3192607

34. Li M, Meng Z, Yu S, Li J, Wang Y, Yang W, Wu H. Baicalein ameliorates cerebral ischemia-reperfusion injury by inhibiting ferroptosis via regulating GPX4/ACSL4/ACSL3 axis. Chem Biol Interact. 2022;366:110137. doi: 10.1016/j.cbi.2022.110137

35. Tuo QZ, Lei P, Jackman KA, Li XL, Xiong H, Li XL, Liuyang ZY, Roisman L, Zhang ST, Ayton S, et al. Tau-mediated iron export prevents ferroptotic damage after ischemic stroke. Mol Psychiatry. 2017;22:1520–1530. doi: 10.1038/mp.2017.171

36. Gaschler MM, Andia AA, Liu H, Csuka JM, Hurlocker B, Vaiana CA, Heindel DW, Zuckerman DS, Bos PH, Reznik E, et al. FINO(2) initiates ferroptosis through GPX4 inactivation and iron oxidation. Nat Chem Biol. 2018;14:507–515. doi: 10.1038/s41589-018-0031-6

37. Bai M, Cui N, Liao Y, Guo C, Li L, Yin Y, Wen A, Wang J, Ye W, Ding Y. Astrocytes and microglia-targeted Danshensu liposomes enhance the therapeutic effects on cerebral ischemia-reperfusion injury. J Control Release. 2023;364:473–489. doi: 10.1016/j.jconrel.2023.11.002

38. Alim I, Caulfield JT, Chen Y, Swarup V, Geschwind DH, Ivanova E, Seravalli J, Ai Y, Sansing LH, Ste.Marie EJ, et al. Selenium Drives a Transcriptional Adaptive Program to Block Ferroptosis and Treat Stroke. Cell. 2019;177:1262–1279.e1225. doi: 10.1016/j.cell.2019.03.032

39. Sanguigno L, Guida N, Anzilotti S, Cuomo O, Mascolo L, Serani A, Brancaccio P, Pennacchio G, Licastro E, Pignataro G, et al. Stroke by inducing HDAC9-dependent deacetylation of HIF-1 and Sp1, promotes TfR1 transcription and GPX4 reduction, thus determining ferroptotic neuronal death. Int J Biol Sci. 2023;19:2695–2710. doi: 10.7150/ijbs.80735

40. Stockwell BR, Friedmann Angeli JP, Bayir H, Bush AI, Conrad M, Dixon SJ, Fulda S, Gascón S, Hatzios SK, Kagan VE, et al. Ferroptosis: A Regulated Cell Death Nexus Linking Metabolism, Redox Biology, and Disease. Cell. 2017;171:273–285. doi: 10.1016/j.cell.2017.09.021

41. Yuan Q, Yuan Y, Zheng Y, Sheng R, Liu L, Xie F, Tan J. Anti-cerebral ischemia reperfusion injury of polysaccharides: A review of the mechanisms. Biomed Pharmacother. 2021;137:111303. doi: 10.1016/j.biopha.2021.111303

42. Jia C, Xiang Z, Zhang P, Liu L, Zhu X, Yu R, Liu Z, Wang S, Liu K, Wang Z, et al. Selenium-SelK-GPX4 axis protects nucleus pulposus cells against mechanical overloading-induced ferroptosis and attenuates senescence of intervertebral disc. Cellular and Molecular Life Sciences. 2024;81. doi: 10.1007/s00018-023-05067-1

43. Chen J, Yang L, Geng L, He J, Chen L, Sun Q, Zhao J, Wang X. Inhibition of Acyl-CoA Synthetase Long-Chain Family Member 4 Facilitates Neurological Recovery After Stroke by Regulation Ferroptosis. Front Cell Neurosci. 2021;15:632354. doi: 10.3389/fncel.2021.632354

